# LRRC55 modulates BK channels to support Purkinje cell plasticity and motor coordination

**DOI:** 10.64898/2026.01.30.702921

**Authors:** Xin Guan, Jiusheng Yan

**Author notes:** Correspondence: Correspondence should be addressed to J.Y.

## Abstract

Large-conductance Ca^2+^- and voltage-activated K^+^ (BK) channels are widely expressed, including in the brain where they shape neuronal excitability. Their physiological functions are strongly influenced by cell-type-specific auxiliary subunits. The auxiliary γ3 subunit (LRRC55) enhances BK-channel activation by shifting voltage-dependent gating toward more negative potentials; however, its protein distribution and in vivo function remain unclear. Here, we generated knock-in mice carrying a C-terminal epitope tag on endogenous LRRC55 to map its expression, and *Lrrc55* knockout mice to test its function. LRRC55 protein was selectively enriched in cerebellar Purkinje cells. *Lrrc55* deletion produced ataxia-like impairments in gait, balance, and coordination. In acute slices, pharmacological BK-channel block with paxilline altered Purkinje cell simple- and complex-spike firing in wild-type mice, whereas these BK-dependent effects were largely absent in *Lrrc55* knockouts, indicating that LRRC55 is required for BK channels to shape Purkinje cell firing under these conditions. Moreover, LRRC55 loss disrupted cerebellar synaptic plasticity, abolishing parallel fiber–Purkinje cell long-term potentiation and eliminating climbing fiber–Purkinje cell long-term depression, phenocopying paxilline in wild-type cells. Together, these results identify LRRC55 as a Purkinje-cell-enriched auxiliary subunit that is essential for BK-dependent excitability and plasticity and that supports normal cerebellar motor function.

## Introduction

Ion channel regulation by auxiliary subunits is a critical mechanism that generates functional diversity and contributes to the variability of electrical signaling across tissues and cell types (1, 2). The big/large-conductance, calcium- and voltage-activated K^+^ (BK) channel is a distinctive member of the potassium channel family, uniquely integrating cellular excitability and calcium signaling through dual regulation by membrane voltage and intracellular Ca^2+^ and by its exceptionally large single-channel conductance (3–5). In central neurons, BK channels mediate the repolarization and fast afterhyperpolarization of action potentials (6, 7), provide negative feedback regulation of Cav channels(8, 9), shape dendritic Ca^2+^ spikes (10), and regulate neurotransmitter release (11–14).

BK channel function is tightly modulated by different auxiliary β and γ subunits, as well as the LINGO1 protein, enabling tissue-specific gating and pharmacological properties (15–18). The 4 BK channel γ subunits are a group of leucine-rich repeat (LRR) containing (LRRC) membrane proteins: LRRC26, LRRC52, LRRC55, and LRRC38, designated as the γ1, γ2, γ3, and γ4 subunits, respectively (19, 20). They all contain an N-terminal signal peptide, an extracellular LRR domain, a single TM segment, and a short intracellular C-terminus (19–21).They possess distinct capabilities in potentiating BK channel activation by shifting the BK channel’s voltage dependence of activation in the hyperpolarizing direction over an exceptionally large range of approximately 145 mV (γ1), 100 mV (γ2), 50 mV (γ3), and 20 mV (γ4) (19, 20). The γ1 subunit is mainly expressed in secretory glands or organs (20) and plays an important role in resting K^+^ efflux and fluid secretion (22, 23). The γ2 subunit is predominantly expressed in the testes (20) and also potently modulates BK channels in cochlear inner hair cells (24).

The BK channel auxiliary γ3 subunit LRRC55 at the mRNA level is predominantly found in the brain (20), with the cerebellum among the few regions showing strong expression by in situ RNA hybridization (25). BK channels are also abundantly expressed in the cerebellum, particularly in Purkinje cells (PCs) (26, 27), where they play a significant role in motor coordination and learning (8, 26, 28, 29). PCs serve as the principal and sole output neurons of the cerebellar cortex, sending GABAergic inhibitory projections to the deep cerebellar nuclei and forming the core of cerebellar circuits. They also receive two types of excitatory inputs: parallel fibers (PFs) from granule cells and climbing fibers (CFs) from the inferior olive, targeting proximal dendrites and dendritic spines, respectively (30–32). Synaptic plasticity at PF-PC and CF-PC synapses modulates PC activity. Notably, both global and PC-specific deletion or dysregulation of BK channels leads to PC dysfunction and cerebellar ataxia in mice (26, 28, 29). Moreover, a loss-of-function BKα mutation in humans causes progressive cerebellar ataxia (33).

Despite its selective expression and potential role in the nervous system, the distribution of LRRC55 at the protein level in the brain and its neurophysiological role remain unknown, largely due to the lack of suitable antibodies and animal models. Understanding how LRRC55 modulates BK channel function in cerebellar Purkinje cells could provide important insights into the molecular mechanisms underlying motor coordination and learning. To address this, we generated knock-in mice carrying a 3×HA-V5 epitope tag at the C terminus of endogenous LRRC55 to map its protein expression, and we created *Lrrc55* knockout mice to investigate its functional role in cerebellar circuits. These animal models provide a platform to dissect the contribution of LRRC55 to BK channel regulation and cerebellar physiology.

## Results

### LRRC55 protein is selectively expressed in cerebellar Purkinje cells

To investigate the role of LRRC55 in brain function, precise assessment of its expression at the protein level is essential, as transcript levels often do not correlate with protein abundance and cannot reveal subcellular localization. However, no specific antibody against LRRC55 is currently available. To overcome this limitation, we generated LRRC55-HA(3×)-V5 knock-in mice on a C57BL/6 background, in which 3×HA (3 tandem HA) and V5 epitope tags were fused to the C-terminus of endogenous LRRC55 (**Fig. 1A & S1A**) using a homology-directed repair (HDR)-based CRISPR/Cas12a genome editing method (34). HA and V5 tags were chosen for their high specificity, as evidenced by the low background observed in immunohistochemical assays of mouse brain tissue (35, 36). Immunofluorescence with anti-HA antibody showed minimal background in WT mouse brain sections (**Fig. 1 B**), confirming antibody specificity. In knock-in mice, HA labeling of endogenous LRRC55 was detected mainly in the olfactory bulb and cerebellum. Within the cerebellum, the HA signal was enriched in Purkinje cell somata and dendritic regions, with the strongest labeling in the Purkinje cell layer (PL) and extending into the molecular layer (ML), whereas the granule cell layer (GL) exhibited low signal. (**Fig. 1 B**). This result of LRRC55 expression at the protein level is consistent with LRRC55 promoter-driven EGFP expression (**Fig. S1B**) in the BAC-transgenic mouse line Tg(LRRC55-EGFP)KS290Gsat (MMRRC #031905-UCD)(37), and as well as with prior in situ hybridization mapping of LRRC55 mRNA in the mouse brain(25).

**Figure 1.**
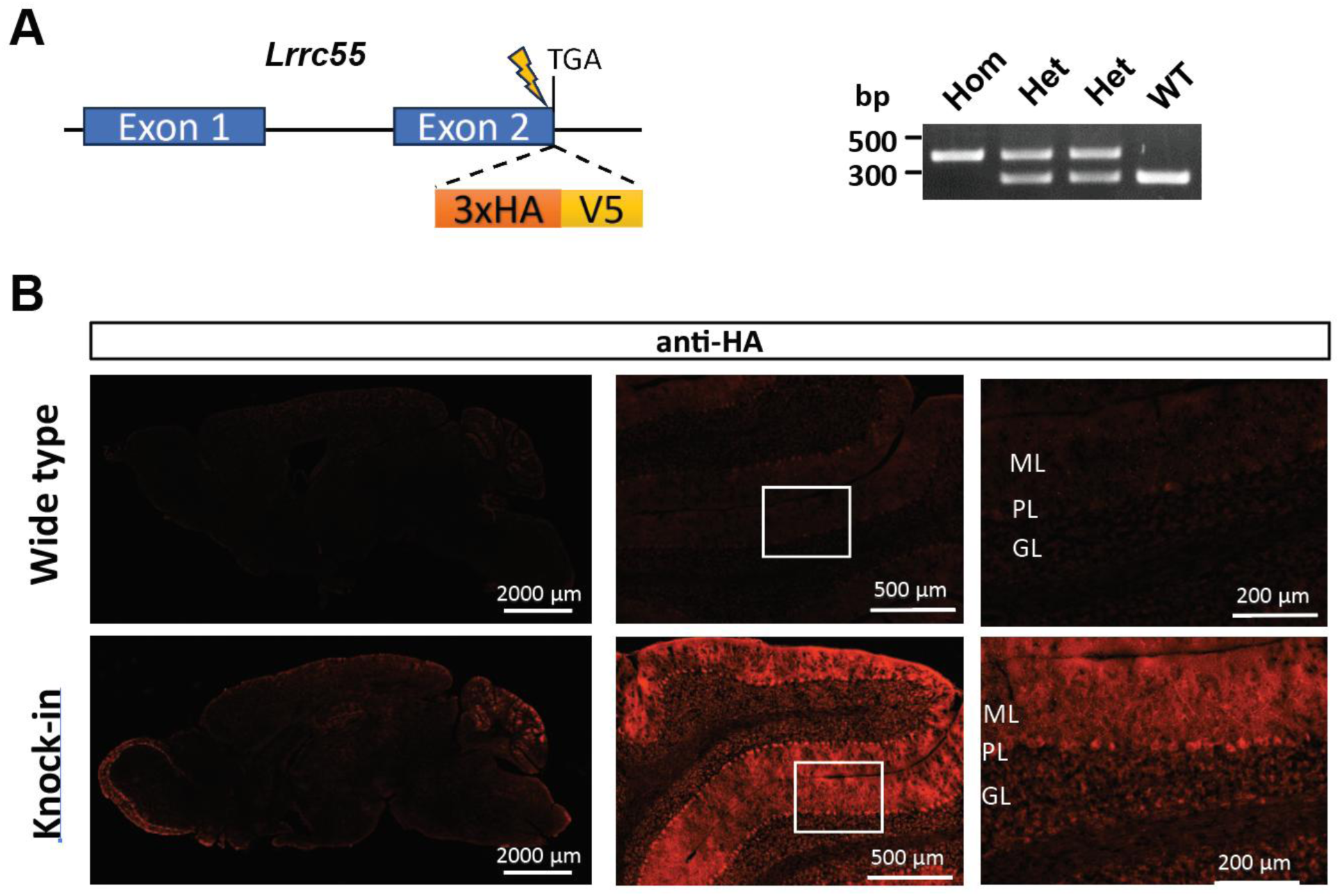
LRRC55 is selectively expressed in cerebellar Purkinje cells. (A) Schematic of the *Lrrc55* knock-in (KI) allele, in which a 3*×*HA–V5 epitope tag is fused to the C terminus of endogenous LRRC55, and representative genotyping PCR for wild-type (WT), heterozygous (Het), and homozygous (Hom) KI mice. (B) Anti-HA immunofluorescence in WT and *Lrrc55*-KI homozygous mouse brain sections showing LRRC55-3*×*HA–V5 signal in the whole brain (left) and cerebellum (middle/right).

### *Lrrc55*-KO mice exhibit impaired motor coordination

To study LRRC55 function in vivo, *Lrrc55* knockout (KO) mice were generated on a C57BL/6 background using the TALENs (transcription activator-like effector nucleases) genome editing method (38, 39), targeting exon 1 of the LRRC55 gene. The *Lrrc55*-KO mice had a 22 bp deletion after the amino acid sequence position 33, resulting in frameshift deletions of nearly the entire LRRC55 protein, except for the short N-terminal signal peptide region (**Fig. 2A & S2A**). The KO mice were backcrossed with WT mice for 6 generations to minimize off-target effects and generic background heterogeneity. Immunofluorescence using an anti-BK antibody revealed no significant difference in BK channel expression in Purkinje cells, granular cells, and neuophi between WT and *Lrrc55*-KO mice, suggesting that LRRC55 has minimal influence on BK channel expression (**Fig. S1C**).

**Figure 2.**
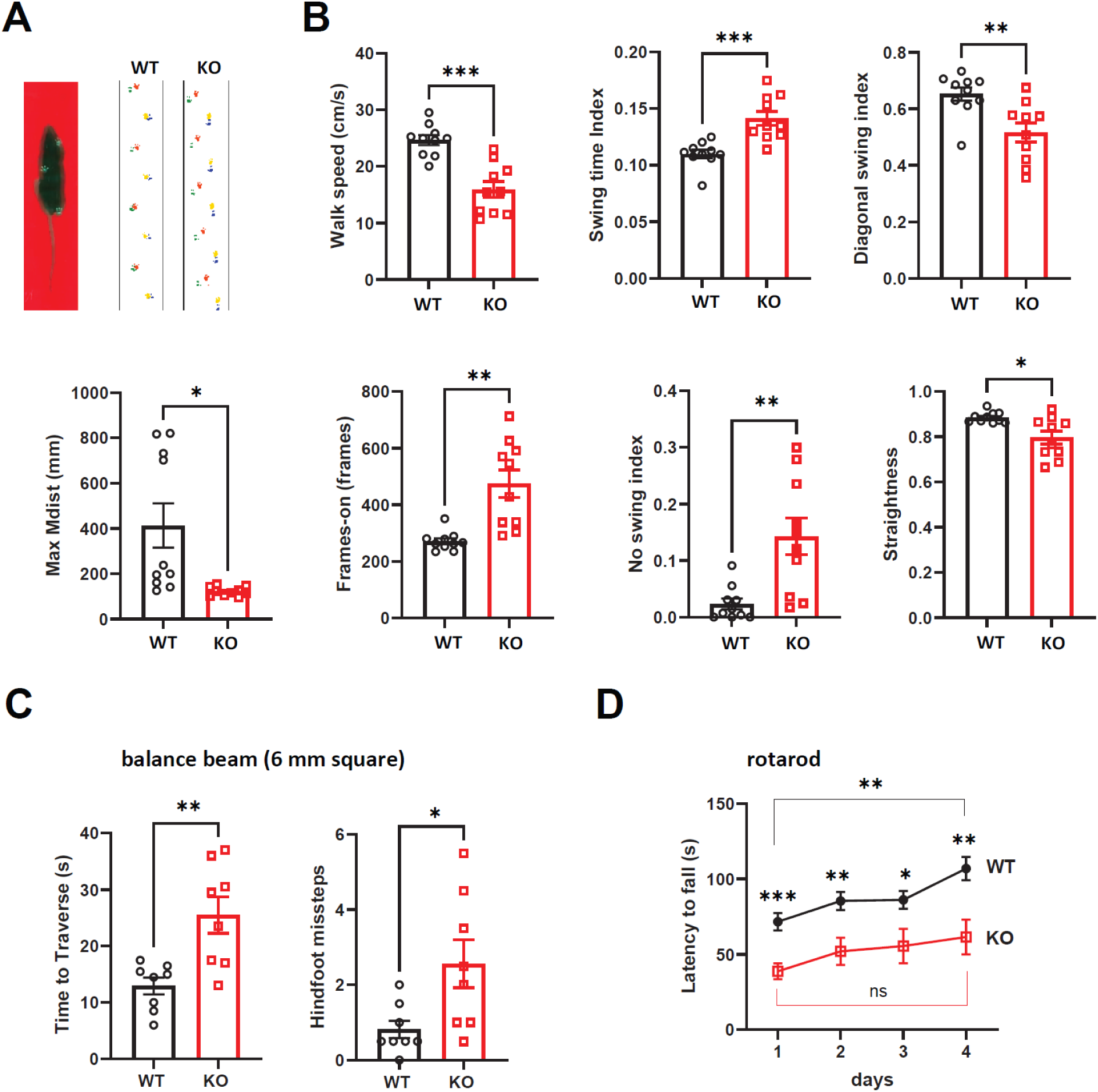
*Lrrc55* knockout mice display ataxia-like motor deficits. (A) Schematic of the optical footprint detection setup and representative gait footprints from WT and *Lrrc55*-KO mice recorded with the MouseWalker system. (B) Automated MouseWalker gait metrics for WT (n =10) and *Lrrc55*-KO (n = 10) mice, including walking speed, swing time index, diagonal swing index, maximum paw excursion distance (Max Mdist), walkway contact (frames-on), fraction of time without swing (no-swing index; all paws on the ground), and walking straightness. (C) Balance beam performance on a 6-mm square beam: traversal time and number of hindlimb slips for WT (n = 8) and *Lrrc55*-KO mice (n = 8). (D) Accelerating rotarod performance (latency to fall) for WT (n = 9) and *Lrrc55*-KO mice (n = 9). Data are mean ± SEM. *, p < 0.05; **, p < 0.01; ***, p < 0.001; ns, not significant.

From open field and hanging wire tests, *Lrrc55*-KO mice showed no significant differences compared with WT controls in locomotor activity, as measured by total travel distance (**Fig. S2B**), or in grip/muscle strength, as measured by latency to fall (**Fig. S2C**). These results indicate that gross motor function was preserved, permitting subsequent assessment of cerebellum-related behaviors in motor coordination and learning. To quantitatively evaluate gait, we employed a sensitive automated MouseWalker system (**Fig. 2A**) following established methods (40, 41). Using this approach, we observed that *Lrrc55*-KO mice exhibited gait abnormalities compared with WT mice (**Fig. 2B**). KO mice walked significantly slower, with prolonged swing times (measured as average swing time index) and a reduced diagonal swing index (**Fig. 2B**). The longer swing phase reflects slowed paw advancement during each step, while the reduced diagonal swing index indicates impaired contralateral limb coupling. In addition, KO mice showed reduced maximum excursion distances of paw movements (Max Mdist), increased walkway contact times (frames-on), and a markedly larger fraction of time without swing (no swing: all four paws simultaneously on the ground) (**Fig. 2B**), reflecting restricted limb excursion and a cautious gait characterized by slower progression, prolonged full-body support, and reduced locomotor dynamism. Moreover, KO mice exhibited less straight paw trajectories, indicating impaired precision of limb control and step guidance during locomotion (**Fig. 2B**).

In the balance beam task, *Lrrc55*-KO mice required significantly more time than WT controls to traverse both the 6-mm square beam (**Fig. 2C**) and the 12-mm round beam (**Fig. S2D**), and they made markedly more hind foot missteps on each beam (**Fig. 2C, S2D**). In the accelerating rotarod test over four consecutive days, *Lrrc55*-KO mice displayed a significantly shorter latency to fall compared with WT mice (**Fig. 2D**). Across all four days, a two-way repeated-measures ANOVA (Genotype × Day) revealed significant main effects of Genotype (F(_1,16_) = 12.7, p < 0.003) and Day (F(_3,48_) = 12.1, p < 0.0001), reflecting significant differences in overall performance between WT and KO mice. Post hoc Tukey’s multiple comparisons test confirmed significant differences in daily performance between WT and KO mice. Across the training period from Day 1 to Day 4, WT groups significantly improved in latency from Day 1 to Day 4 by 35 s, whereas KO mice exhibited an overall smaller increase (23 s; not statistically significant) in latency. The apparent difference in improvement between WT and KO mice, however, did not reach statistical significance, as reflected by the absence of a significant Genotype × Day interaction (F(_3,48_) = 0.95, p = 0.40).

Collectively, these findings demonstrate that *Lrrc55*-KO mice exhibit an ataxia-like phenotype characterized by impaired gait efficiency, disrupted interlimb coordination, reduced precision of paw placement, and deficits in balance and motor learning, supporting a critical role for LRRC55 in cerebellar motor control.

### LRRC55 is required for the BK channel-mediated modulation of PC firing activities

Cerebellar dysfunction is a primary cause of ataxia, which is frequently associated with impaired PC function (42). PCs are characterized by two distinct firing patterns: spontaneous tonic activity in the form of simple spikes (8, 43, 44), and climbing fiber (CF)-evoked repetitive bursting known as complex spikes (26, 28, 45). BK channels were reported to modulate both types of firing patterns (28, 46). To investigate PC firing dynamics, we performed whole-cell current-clamp recordings in cerebellar slices, in the absence and presence of paxilline, a specific blocker of BK channels that, unlike another similarly commonly used BK-specific blocker, iberiotoxin, is not affected by the brain-specific β4 subunit, thereby providing a more precise measure of BK channel contribution.

In WT mice, unstimulated PCs exhibited spontaneous tonic firing. Application of paxilline significantly increased the average firing frequency from 29.6 ± 3.0 Hz to 49.5 ± 7.4 Hz (n = 17 measurements from 5 mice) (**Fig. 3A, B**). In contrast, the firing frequency of PCs from Lrrc55 knockout (KO) mice remained unchanged before (37.3 ± 3.4 Hz) and after paxilline treatment (35.3 ± 3.9 Hz; n = 16 from 5 mice) (**Fig. 3A, B**). Upon CF stimulation, complex spikes were recorded. Consistent with previous reports on the effect of BK channel deletion (28), paxilline induced an additional spikelet in WT PCs, increasing the average spikelet number from 2.7 to 3.7 before and after BK channel blockade, respectively (n = 7 from 4 mice). (**Fig. 3C, D**). However, in *Lrrc55*-KO PCs, no change in spikelet number was observed in most recordings (6 out of 7 cells), with the average spikelet count remaining essentially unchanged at 2.9 before and 3.0 after paxilline application (n = 7 from 3 mice). (**Fig. 3C, D**). Taken together, these findings demonstrate that LRRC55 is essential for BK channel-mediated modulation of PC firing. In its absence, the influence of BK channels on both spontaneous tonic activity and CF-evoked complex spikes is abolished. It is of note that the recorded simple spike frequency and the number of spikelets in evoked complex spikes were not statistically different between WT and *Lrrc55*-KO PCs, suggesting that some compensatory mechanisms may have arisen in *Lrrc55*-KO mice to counteract the loss of BK channel modulation.

**Figure 3.**
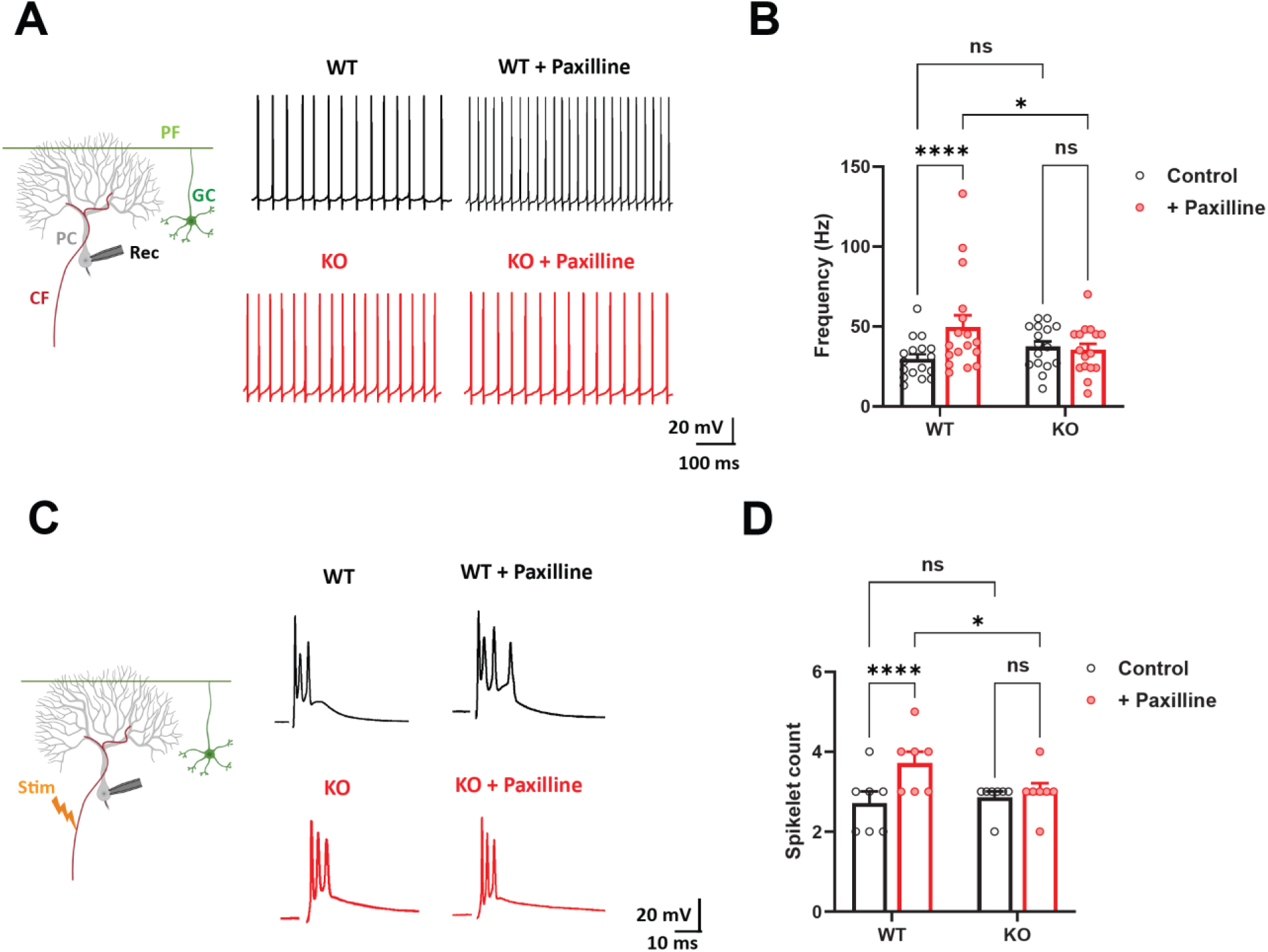
BK-channel blockade alters Purkinje cell firing in WT but not *Lrrc55*-KO mice. (A) Representative spontaneous firing traces from WT (top, black) and *Lrrc55*-KO (bottom, red) PCs before and after paxilline (10 *μ*M). (B) Quantification of spontaneous firing rate before and after application of paxilline in WT (n = 17 cells from 5 mice) and *Lrrc55*-KO mice (n = 16 cells from 5 mice). (C) Representative complex spike recordings from WT (top, black) and *Lrrc55*-KO (bottom, red) Purkinje cells (PCs) before and after application of paxilline (10 *μ*M) during climbing fiber stimulation. (D) Quantification of complex-spike spikelet number before and after application of paxilline in WT (n = 7 cells from 4 mice) and *Lrrc55*-KO mice (n = 7 cells from 3 mice). Schematic cartoons of stimulation/recording configurations are shown at left in (A) and (C). PC, Purkinje cell; GC, granule cell; PF, parallel fiber; CF, climbing fiber; stim, stimulation; Rec, recording electrode. Data are mean ± SEM. *, p < 0.05; **, p < 0.01; ns, not significant.

### LRRC55’s regulation of BK channels is required for long-term potentiation at parallel fiber-Purkinje cell synapses

Parallel fiber–Purkinje cell (PF-PC) synaptic plasticity, expressed as long-term potentiation (LTP) and long-term depression (LTD), plays a central role in cerebellar function by adjusting the strength of excitatory input onto Purkinje cells, thereby shaping motor learning, coordination, and adaptive control of movement (47). To assess the role of LRRC55 in PF–PC synaptic plasticity, EPSCs (excitatory postsynaptic currents) of PCs were recorded in cerebellar slices using whole-cell voltage-clamp recordings combined with parallel fiber (PF) stimulation, as previously described (48). In WT mice, PF stimulation at 1 Hz for 5 min reliably induced PF–PC LTP, as indicated by a gradual increase in EPSC amplitude to 150% ± 12% of baseline (n = 6 from 5 mice) within 15 min after stimulation. In contrast, the magnitude of LTP was largely abolished in *Lrrc55*-KO mice, with EPSCs reaching only 113% ± 5% of baseline (n = 9 from 5 mice), and the residual potentiation appeared with a significant delay (**Fig. 4A, B, C**). Furthermore, paxilline application completely blocked LTP at PF–PC synapses in WT mice, leaving EPSC amplitudes unchanged at 103% ± 11% of baseline (n = 10 from 5 mice) (**Fig. 4A, B, C**). Because LTP was already largely absent in *Lrrc55*-KO mice, paxilline was not applied in KO slices, as additional BK channel blockade would not be expected to produce further measurable suppression. These results indicate that both LRRC55 and BK channels were required for PF-PC LTP.

**Figure 4.**
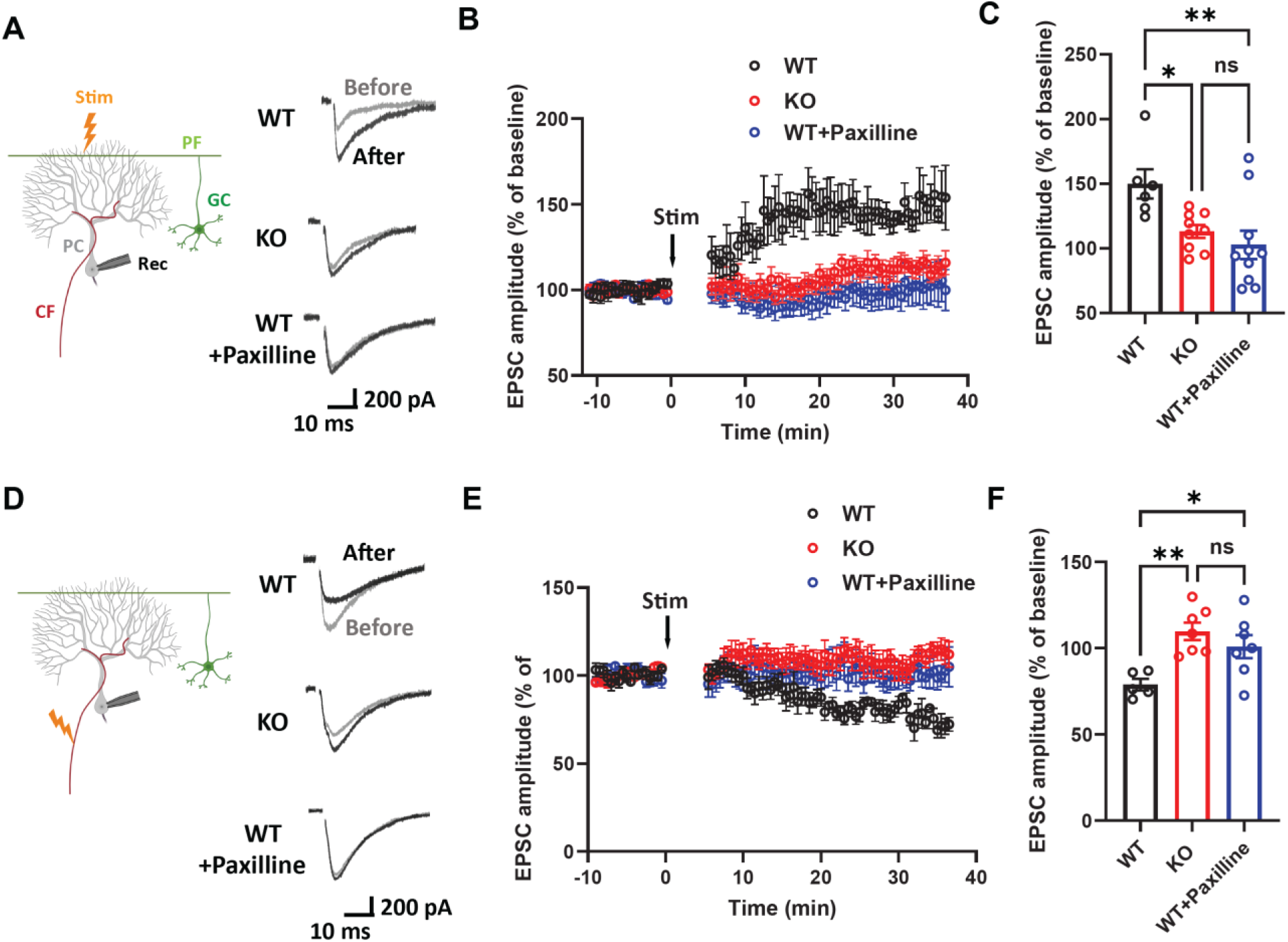
LRRC55 is required for BK-dependent synaptic plasticity in Purkinje cells. (A–C) Parallel fiber (PF)–PC long-term potentiation (LTP) is impaired in *Lrrc55*-KO slices and in WT slices treated with paxilline, compared with WT controls (WT: n = 6 cells from 5 mice; *Lrrc55*-KO: n = 9 cells from 5 mice; WT + paxilline: n = 10 cells from 5 mice). (A) Representative PF-evoked EPSC traces before (gray) and after (black) LTP induction. (B) Time course of normalized EPSC amplitudes. (C) Mean normalized EPSC amplitude measured 30–35 min after induction. (D–F) Climbing fiber (CF)–PC long-term depression (LTD) is induced in WT slices but not in *Lrrc55*-KO slices or WT slices treated with paxilline (WT: n = 7 cells from 5 mice; *Lrrc55*-KO: n = 7 cells from 5 mice; WT + paxilline: n = 7 cells from 5 mice). (D) Representative CF-evoked EPSC traces before (gray) and after (black) LTD induction. (E) Time course of normalized EPSC amplitudes. (F) Mean normalized EPSC amplitude measured 30–35 min after induction. Schematic cartoons of stimulation/recording configurations are shown at left in (A) and (D). Data are mean ± SEM. *, p < 0.05; **, p < 0.01; ns, not significant.

We also examined long-term depression (LTD) at PF–PC synapses using a conjunctive induction protocol, consisting of parallel fiber burst stimulation (300 pulses at 1 Hz) paired with somatic depolarization of Purkinje cells to 0 mV for 200 ms, as previously described (49). In both WT and *Lrrc55*-KO mice, this stimulation reliably induced LTD, reflected by a sustained reduction in EPSC amplitude (**Fig. S3A, B, C**). The extent of LTD was comparable between genotypes, with EPSCs reduced to 62% ± 9% of baseline in WT mice (n = 6 from 5 mice) and 68% ± 3% in *Lrrc55*-KO mice (n = 9 from 5 mice). Furthermore, treatment with paxilline did not significantly alter LTD in WT mice, as EPSCs were reduced to 71% ± 6% of baseline (n = 10 from 5 mice). Because LTD was unaffected in Lrrc55-KO mice, paxilline was not tested in KO slices, as additional BK channel blockade would not be expected to produce further effects. These results indicate that, unlike LTP, LTD at PF–PC synapses is independent on LRRC55 or BK channel activity.

To evaluate presynaptic function at PF–PC synapses, we measured paired-pulse facilitation (PPF), a form of short-term plasticity that reflects changes in presynaptic release probability (50). Both WT and *Lrrc55*-KO mice exhibited comparable levels of PPF across a range of interpulse intervals (20–1000 ms), with no significant differences detected between the two groups (WT, n = 7 from 4 mice; *Lrrc55*-KO, n = 7 from 4 mice) (**Fig. S3D, E)**. These results indicate that deletion of *Lrrc55* does not affect presynaptic neurotransmitter release at PF–PC synapses, consistent with the relatively low expression of LRRC55 in the granule cell layer compared with Purkinje cells and the molecular layer (**Fig. 1B**).

### LRRC55’s regulation of BK channels is required for long-term depression at climbing fiber-Purkinje cell synapses

Climbing fiber activity conveys powerful error signals essential for motor learning tasks (51), and LTD at CF–PC synapses has been proposed to fine-tune the strength of this error-teaching signal (32). Previous studies showed that CF–PC LTD can be completely blocked by iberiotoxin (29), implicating BK channels in its induction. To investigate the role of LRRC55 in this form of plasticity, we induced CF–PC LTD by depolarizing (–65 mV to –55 mV) PCs in current-clamp mode, followed by tetanic climbing fiber stimulation at 5 Hz for 30 s, using the same protocol as previously established (32). In WT mice, this protocol reliably induced CF–PC LTD, as reflected by a reduction in CF–EPSC amplitude to 79% ± 3% of baseline within 30 min after tetanic stimulation (n = 5 cells from 5 mice) (**Fig. 4C, D, E**). In contrast, the same stimulation failed to induce LTD in *Lrrc55*-KO mice or in WT mice treated with paxilline, with CF–EPSC amplitudes remaining at 110% ± 5% and 101% ± 7% of baseline, respectively (n = 7 cells from 5 mice for each group; **Fig. 4C, D, E**). To assess presynaptic function, we measured paired-pulse depression (PPD) at CF–PC synapses and found no difference between *Lrrc55*-KO (n = 7 from 4 mice) and WT mice (n = 6 from 4 mice) (**Fig. S3F, G**), indicating that presynaptic release was unaffected. Together, these findings demonstrate that CF–PC LTD requires not only BK channel activity but also its regulation by LRRC55.

## Discussion

BK channels are prominently expressed throughout the Purkinje cell (PC) compartments, including soma, dendrites, and axon (26, 27), and extensive genetic and pharmacological work has established that perturbing BK function is sufficient to disrupt PC output and produce cerebellar ataxia-like phenotypes (26, 28, 29, 33). However, it remains unclear how BK channels are enabled in cerebellar function, particularly which endogenous PC-enriched factor confers this capability. Our findings place LRRC55 (the BK *γ*3 subunit) as a key component of this tuning in cerebellar PCs. Specifically, LRRC55 is selectively enriched in PCs, and its deletion causes ataxia-like behaviors, removes BK-dependent modulation of PC firing, and disrupts two forms of cerebellar synaptic plasticity (PF–PC LTP and CF–PC LTD).

### LRRC55 as a Purkinje cell-enriched BK channel auxiliary subunit

A technical barrier in mapping LRRC55 expression at the protein level has been the lack of suitable antibody or reagents. Using an epitope-tagged knock-in allele, we detect robust LRRC55 signal in the Purkinje cell layer and molecular layer, consistent with somatodendritic enrichment. This restricted distribution contrasts with the broader expression of BK *α* and with auxiliary subunits such as *β*4, supporting the idea that BK channels in PCs assemble with a distinct auxiliary-subunit complement. Importantly, our data does not indicate a major change in BK protein expression in *Lrrc55*-KO cerebellum, arguing that the phenotype is more likely explained by altered BK channel function (gating properties) rather than by a loss of channel expression.

### LRRC55 is indispensable for BK channel function in Purkinje cells

In WT PCs, paxilline increases simple-spike firing rate and increases complex-spike spikelet number, consistent with prior work showing that BK currents shape repolarization, afterhyperpolarization, and complex spike waveform (7, 28, 46). In contrast, these modulatory effects of BK channels were abolished in *Lrrc55*-KO PCs, demonstrating that LRRC55 is required to couple BK channel activity to intrinsic excitability. Unlike another brain-specific auxiliary subunit β4, which dampens BK activity (52), LRRC55 potentiates BK channel activation (20). One might therefore predict that LRRC55 simply increases the magnitude of BK-channel contributions to Purkinje cell (PC) function. Instead, LRRC55 loss effectively uncouples BK channels from excitability control, indicating that LRRC55 is not merely an amplifier but a critical determinant that enables BK channels to regulate PC excitability. Baseline firing frequency and spikelet number were comparable between genotypes, suggesting that LRRC55 deletion may induce compensatory ionic conductances that help stabilize firing. However, such compensation cannot replace the dynamic modulation normally provided by LRRC55-modulated BK channels, and LRRC55 deletion nevertheless produced behavioral ataxia that parallels key aspects of BK channel loss-of-function models (26, 28, 29, 33). Thus, LRRC55 is an essential part of the molecular machinery that makes BK channels functionally relevant in PCs.

### LRRC55 contributes to cerebellar plasticity and motor coordination

A key advance of this work is the identification of LRRC55–BK channel signaling as essential for long-term plasticity at both PF and CF synapses. PF–PC LTP and CF–PC LTD were abolished in *Lrrc55*-KO slices and in WT slices treated with paxilline, whereas PF–PC LTD and presynaptic paired-pulse facilitation or depression were unchanged. Our findings thus reveal LRRC55–BK channel signaling as a shared requirement for both potentiation of PF inputs and depression of CF inputs. Prior work demonstrated that climbing fiber LTD requires postsynaptic Ca²⁺ (32) and is blocked by BK channel inhibitors (29). Because BK channels shape Ca²⁺-dependent excitability and repolarization in Purkinje cells (7, 9), the simultaneous loss of PF–PC LTP and CF–PC LTD in *Lrrc55*-KO mice likely arise from a common disruption of the related postsynaptic Ca²⁺-dependent signaling (24, 38). At the circuit level, PCs must integrate dense PF input with instructive CF signals to maintain timing precision and to support adaptive motor learning. The *Lrrc55* ablation-induced postsynaptic deficits is likely to impair the adaptive tuning of PC output within the olivo-cerebellar loop. Our behavioral data, combined with preserved gross locomotion and strength measures, are consistent with a primary cerebellar contribution for the ataxia-like behaviors in *Lrrc55*-KO mice. The combined loss of BK-dependent modulation of firing and the elimination of PF–PC LTP and CF–PC LTD provides a mechanistically informed explanation for impaired coordination and motor learning in the absence of LRRC55.

### Conclusion and limitation

Our findings underscore the importance of an auxiliary subunit in defining the physiological roles of an ion channel in specific neuron and circuit. It is of note that because the KO is constitutive, developmental compensation and its loss in some other brain region may complicate the results or mask additional roles of LRRC55. Despite this limitation, the convergence of selective LRRC55 expression in PCs, loss of paxilline-sensitive modulation of firing, and a selective plasticity phenotype supports a model in which LRRC55 is an essential in vivo determinant of BK-dependent Purkinje cell excitability and plasticity, with potential relevance to cerebellar ataxia.

## Methods and Materials

### Mice

To generate the LRRC55 knock-in mice, we tagged performed CRISPR-assisted targeting endogenous LRRC55 with a 3×HA-V5 epitope in C57BL/6J embryonic stem cells. LRRC55 knockout mice were generated using the TALENs (transcription activator-like effector nucleases) genome editing method (38, 39), targeting exon 1 of the LRRC55 gene. The *Lrrc55*-KO mice contain 22 bp deletions after the corresponding amino acid sequence positions 33. We had backcrossed the mice with C57BL/6 mice for 6 generations to minimize off-target effects and generic background heterogeneity at the beginning of this study. All animal experiments were carried out according to protocols and guidelines approved by the Institutional Animal Care and Use Committee of The University of Texas MD Anderson Cancer Center.

### Behavioral Tests

All mice used for behavioral testing were age-matched across groups (8–12 weeks old; both sexes included).

### Footprint analysis

A sensitive automated gait analysis MouseWalker system by constructing a frustrated total internal reflection (fTIR)-based optical footprint detection apparatus were used following previously described methods (40). The mouse’s paw contact with the glass surface generates light reflection in the form of illuminated footprints, which are recorded with a high speed (≥100fps) video camera and analyzed using a MATLAB software tool MouseWalker (40, 41).

### Balance beam test

Mice were trained to walk from the starting point along a 1-m long, 12-mm in diameter round or 6-mm wide square beam placed 50 cm above bedding. Mice were allowed to stay on the beam for 60s maximum. The latency to traverse each beam and the slip number of the hind feet were recorded for each trail.

### Accelerating rotarod test

Mice were placed on the rod of rotating rod apparatus (Harvard Apparatus), and the rod was accelerated from 4 to 40 rpm in 5 min. latency to fall off were recorded every day for 4 days.

### Open field test

Mice were placed in the center of a 40 x 40 cm open field box and were allowed to move freely for 1 h. The total distance moved was recorded and analyzed with an automated video-tracking system (EthoVision XT, Noldus Information Technology).

### Hanging wire test

Mice were placed on a wire cage lid and the lid was turned upside down at 20 cm height above the cage to prevent the animal from climbing down. The time elapsed before falling off was recorded with the maximum time at 60s.

#### Immunofluorescence

Mice were anesthetized and perfused transcardially with 4% paraformaldehyde in PBS, and their brains were postfixed in the same fixative buffer overnight at 4 °C. Then brains were moved to 15% and 30% sucrose solutions (in PBS) for cryoprotection. Brains were embedded in Tissue Tek OCT medium (Sakura) and cryosectioned at 20 µm. The sections were then processed for immunolabeling using standard procedures. Rabbit anti-HA (1:500; Cell Signaling Technology), mouse anti-GFP (1:150, Santa Cruz), and rabbit anti-BK (1:100; Alomone labs) were diluted in immunoreaction enhancer solution (“Can Get Signal”, Toyobo) and used as primary antibody. Nuclei were stained with DAPI (Sigma). Images were captured using a BioTek Cytation 5 cell imaging multimode reader (Agilent) and an Andor Revolution XDi WD Spinning Disk Confocal microscope.

#### Electrophysiology

Cerebellum slices (250 μm) were prepared from 3- to 4-wk-old mice (both sexes included) by using a vibratome (Leica Biosystems) and kept at 37 °C for 0.5 h and then at room temperature in artificial cerebral spinal fluid (ACSF) containing 124 mM NaCl, 5 mM KCl, 1.25 mM Na_2_HPO_4_, 2 mM MgSO_4_, 2mM CaCl_2_, 26 mM NaHCO_3_, and 25 mM D-glucose aerated with 95% O_2_ and 5% CO_2_. The cerebellum was visualized under an infrared differential interference contrast optics microscope (Zeiss). Whole-cell patch clamp recordings were performed on cerebellar Purkinje cells with a MultiClamp 700B amplifier (Axon Instruments) at 31–33 °C. Pipette electrodes (3–5 MΩ) were filled with 145 mM K-gluconate, 5 mM NaCl, 1 mM MgCl2, 2 mM Mg·ATP, 0.1 mM Na·GTP, 0.2 mM EGTA, and 10 mM Hepes (pH 7.2). Only neurons with a resting membrane potential less than or equal to −65 mV and a stable series resistance or capacitance were used for data recording. All experiments were performed in the presence of picrotoxin (100 μM) to reduce GABAergic contributions. A bipolar stimulating electrode was placed in the molecular layer or granule cell layer to stimulate the parallel fibers (PFs) or climbing fibers (CFs), respectively, by using a Master-9 pulse stimulator (A.M.P.I.). PFs were stimulated in the molecular layer at about two-thirds of the distance between the PC layer and the pial surface. CFs were stimulated though a bipolar electrode placed in the granule cell layer and moved around until an all-or-none response was evoked. For Purkinje cells spontaneous action potential recording and complex spike elicited by stimulating climbing fiber recording, whole cell current-clamp mode was performed on Purkinje cells. For PF-PC LTP recording, PFs was stimulated at the test responses of 1 Hz for 5 min (48). For PF-PC LTD recording, PFs was stimulated at the test responses of 300 times at 1 Hz, paired with PC depolarization (0 mV, 200ms) (49). For CF-PC LTD recording, PCs recording switched to current-clamp mode, and small DC currents injected to hold the cell to the range of - 65 to -55 mV, and then CFs were stimulated at the test responses of 5 Hz for 30s (32). Test responses were elicited at 0.03 Hz, and four traces of subthreshold EPSCs were averaged. Paxilline (SANTA CRUZ) and picrotoxin (Tocris Bioscience) were dissolved in dimethyl sulfoxide to create stock solutions and finally diluted with ACSF to specific concentrations before each experiment.

## Data Analysis and Statistics

Results are shown as mean ± SEM. For behavioral analysis, Student’s *t* test was used for comparison of two groups. One-way and two-way ANOVA followed by Tukey’s multiple comparisons test were used for comparisons with multiple groups or conditions. P values < 0.05 were considered significant. All statistical analyses were conducted using GraphPad Prism version 10.

## Acknowledgments

This work was supported in part by National Institutes of Health grants NS078152 (to J. Yan).

## Author contributions

X. Guan and J. Yan designed experiments, analyzed data, and wrote the manuscript. X. Guan performed experiments.

## Supplemental materials

**Figure S1.**
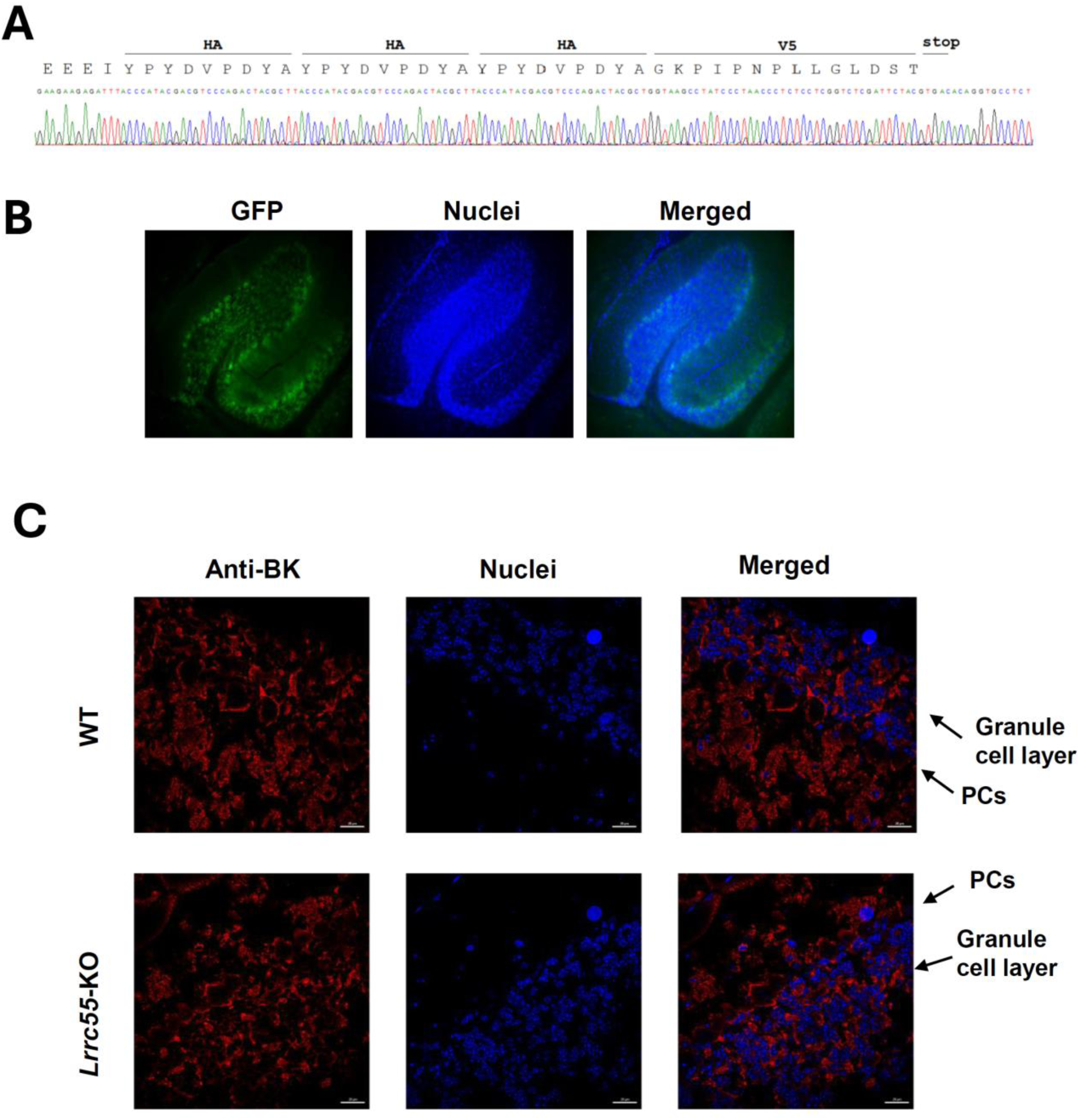
LRRC55 tagging, promoter-driven reporter expression, and BK channel staining. (A) Peptide and DNA sequences encoding the 3*×*HA–V5 tag fused to the C terminus of LRRC55 in *Lrrc55*-KI mice. (B) Immunofluorescence of GFP in brains of the BAC-transgenic mouse line Tg(LRRC55-EGFP)KS290Gsat (MMRRC #031905-UCD), in which GFP expression is driven by the *Lrrc55* promoter. (C) Immunofluorescence for the BK channel *α* subunit in cerebellum from WT and *Lrrc55*-KO mice.

**Figure S2.**
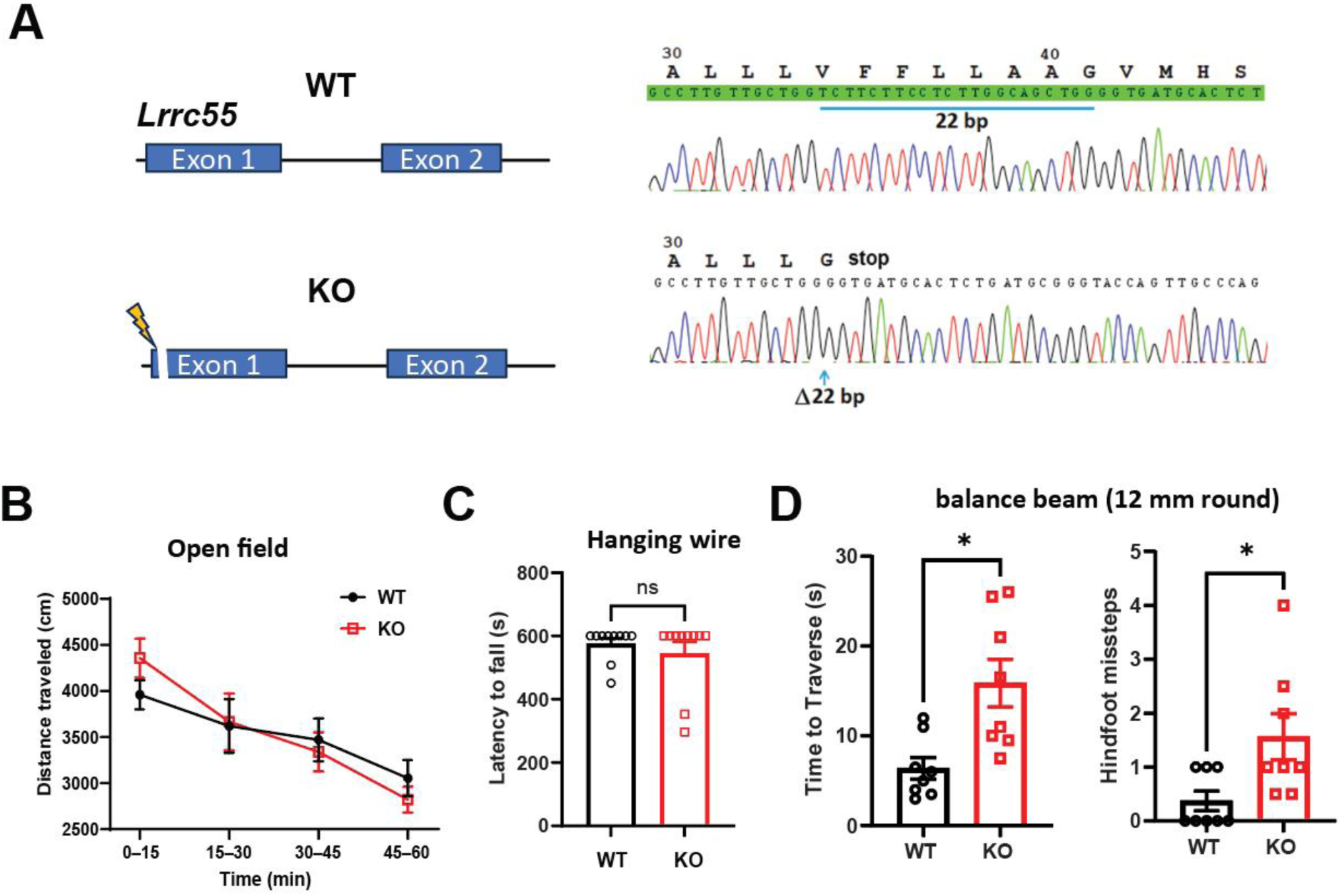
Additional genotyping and behavioral tests. (A) Schematic of the *Lrrc55* locus in WT and KO mice and validation of the KO allele by Sanger sequencing. The KO allele carries a 22-bp deletion in the coding region, resulting in a frameshift early in the coding sequence, leading to a premature stop codon and predicted truncation of the LRRC55 protein. Representative sequencing chromatograms and corresponding translated amino acid sequences are shown for WT and KO alleles. (B) Open field test: distance traveled (cm) in 15-min intervals during a 60-min test by Lrrc55-KO mice (n = 6) and WT mice (n = 4). (C) Hanging wire test: latency to fall (muscle strength) for WT (n = 10) and *Lrrc55*-KO mice (n = 10). (D) Balance beam performance on a 12-mm round beam: traversal time and number of hindlimb slips for WT (n = 8) and *Lrrc55*-KO mice (n = 8). Data are mean ± SEM. *, p < 0.05; ns, not significant.

**Figure S3.**
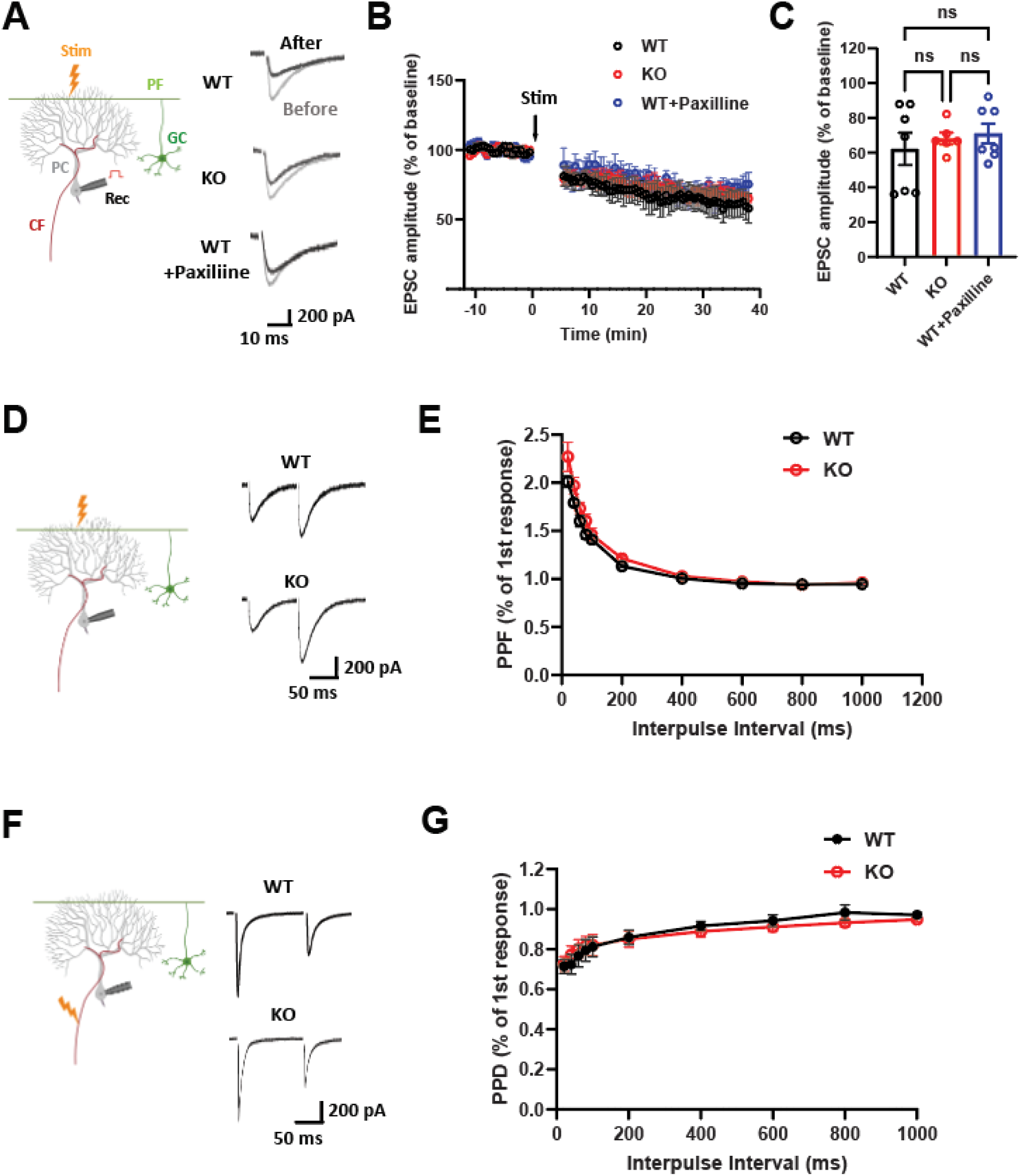
Parallel fiber–Purkinje cell long-term depression and presynaptic short-term plasticity are preserved in Lrrc55-KO mice. (A–C) PF–PC long-term depression (LTD) is comparable between WT and Lrrc55-KO mice and is not affected by paxilline in WT slices (WT: n = 7 cells from 5 mice; Lrrc55-KO: n = 6 cells from 5 mice; WT + paxilline: n = 7 cells from 5 mice). (A) Representative PF-evoked EPSC traces recorded before (gray) and after (black) LTD induction. (B) Time course of normalized EPSC amplitudes.(C) Quantification of normalized EPSC amplitude measured 30–35 min after LTD induction. (D, E) Presynaptic short-term plasticity at PF synapses. (D) Representative paired-pulse EPSC traces from WT and Lrrc55-KO mice. (E) Summary plot of paired-pulse facilitation (PPF) across inter-pulse intervals (WT: n = 7 cells from 4 mice; Lrrc55-KO: n = 7 cells from 4 mice). (F, G) Presynaptic short-term plasticity at CF synapses. (F) Representative paired-pulse EPSC traces from WT and Lrrc55-KO mice. (G) Summary plot of paired-pulse depression (PPD) across inter-pulse intervals (WT: n = 6 cells from 4 mice; Lrrc55-KO: n = 7 cells from 4 mice). Schematic diagrams of stimulation and recording configurations are shown to the left of panels A, D, and F. Data are presented as mean ± SEM. ns, not significant.

## Notes

### Competing Interest Statement

The authors have declared no competing interest.

### Summary of Updates

Minor revision to correct for figure citation, labeling, and legends. Deletion of significance statement.

